# Transcriptome analyses describe the consequences of persistent HIF-1 over-activation in *Caenorhabditis elegans*

**DOI:** 10.1101/2023.11.15.567311

**Authors:** Dingxia Feng, Long Qu

## Abstract

Metazoan animals rely on oxygen for survival, but during normal development and homeostasis, animals are often challenged by hypoxia (low oxygen). In metazoans, many of the critical hypoxia responses are mediated by the evolutionarily conserved hypoxia-inducible transcription factors (HIFs). The stability and activity of HIF complexes are strictly regulated. In the model organism *C. elegans*, HIF-1 stability and activity are negatively regulated by VHL-1, EGL-9, RHY-1 and SWAN-1. Importantly, *C. elegans* mutants carrying strong loss-of-function mutations in these genes are viable, and this provides opportunities to interrogate the molecular consequences of persistent HIF-1 over-activation. We find that the genome-wide gene expression patterns are compellingly similar in these mutants, supporting models in which RHY-1, SWAN-1 and EGL-9 function in common pathway(s) to regulate HIF-1 activity. These studies illuminate the diversified biological roles played by HIF-1, including metabolism, hypoxia and other stress responses, reproduction and development. Genes regulated by persistent HIF-1 over-activation overlap with genes responsive to pathogens, and they overlap with genes regulated by DAF-16. As crucial stress regulators, HIF-1 and DAF-16 converge on key stress-responsive genes and function synergistically to enable hypoxia survival.

## Introduction

As the electron acceptor during oxidative phosphorylation for energy production, oxygen is vital to all the aerobic organisms. Insufficient oxygen availability not only decreases energy production but also changes the redox environment for cellular biochemical reactions [1, 2]. During normal development and disease states, organisms are often challenged by hypoxia. In metazoans, the hypoxia-inducible transcription factors HIFs have evolutionarily conserved roles in mediating critical transcriptional responses to hypoxia. In mammals, HIF complexes regulate genes functioning in angiogenesis, erythropoiesis and glycolysis to assist hypoxia survival [3–6]. HIF’s function and regulation have the potential therapeutic significance in hypoxia-related diseases, such as cancer and stroke. HIF complexes are heterodimers composed of an α subunit and a β subunit, and both subunits are bHLH (basic-helix-loop-helix)-PAS (PER/ARNT/SIM) domain proteins [7]. The human genome encodes three HIFα subunits: HIF-1, -2 and -3α, and three HIFβ/ARNT subunits: HIF-1, -2 and -3β [8]. While HIFβ has multiple bHLH-PAS dimerization partners and is relatively stable and abundant, HIFα is short-lived under well-oxygenated conditions. Thus, HIFα is dedicated to regulate the expression of oxygen-sensitive genes [9–11]. The oxygen-dependent HIFα degradation pathway is conserved and well established. When oxygen levels are high, prolyl hydroxylase domain proteins (PHDs) hydroxylate specific proline residues on HIFα, using oxygen as co-substrate. The hydroxylated HIFα is targeted for proteasomal degradation by an E3 ubiquitin ligase containing the tumor suppressor von Hippel-Lindau (VHL) [2, 10].

HIF and the regulatory network that regulates oxygen-sensitive degradation are conserved and expressed in widely divergent metazoans [12]. HIF and its regulatory system are simplified in the nematode *C. elegans*, and this provides opportunities to interrogate this pathway through genetic analyses [13]. In *C. elegans*, the single counterparts for HIFα and HIFβ are called HIF-1 and AHA-1, respectively [14–16]. While the *hif-1*α -/- mouse dies by E9.0 with severe vascular defects [17, 18], *C. elegans hif-1*(*ia04*) loss-of-function mutants survive and develop normally in room air, although they fail to adapt to hypoxia (0.5% or 1% ambient O_2_) [15, 19, 20]. In *C. elegans*, HIF-1 plays diverse biological roles. In addition to regulating hypoxia response, it regulates responses to other stressors, including heat and toxic chemicals (heavy metal cadmium, ethidium bromide, selenium, nanopolystyrene, silver nanoparticles, tunicamycin, tert-butyl hydroperoxide, hydrogen sulfide and hydrogen cyanide), as well as pathogens (*Staphylococcus aureus*, *Vibrio alginolyticus*, enteropathogenic *Escherichia coli*, and *Pseudomonas aeruginosa* PA14 and PAO1) [21–41]. HIF also contributes to the homeostasis of protein and iron [36, 42–47], reproduction, development and apoptosis [39, 48–50], and neural function and animal behaviors [41, 51–60]. Even more interestingly, HIF-1 has been shown to have roles in regulating animal lifespan [43, 44, 61–77].

The PHD-VHL system for HIF stability regulation is conserved and simplified in *C. elegans*, too. While humans have three PHDs, *C. elegans* has only one counterpart encoded by *egl-9*; and the single VHL homolog in *C. elegans* is encoded by *vhl-1* [14].

The two HIF-1 negative regulators: *rhy-1* and *swan-1* have been shown to regulate the expression of some HIF-1 targets [39, 50]. *rhy-*1 is a multi-pass transmembrane protein [50], and *swan-1* is a conserved WD repeat scaffold protein [78]. *rhy-1(ok1402)* and *swan-1(ok267)* loss-of-function mutations do not dramatically alter HIF-1 protein levels, but increase expression of some of the genes regulated by HIF-1 [39, 50]. Prior studies [14, 39, 50, 79] support a model in which VHL-1 inhibits HIF-1 stability, EGL-9 inhibits HIF-1 stability and activity, RHY-1 and SWAN-1 suppress HIF-1 activity. Loss-of-function mutations in *egl-9* or *rhy-1* and *swan-1;vhl-1* double mutations cause HIF-1 over-activation. We propose that EGL-9, SWAN-1 and RHY-1 function in common pathway(s) to inhibit HIF-1 activity, and EGL-9 and SWAN-1 may form a complex. This model has been further validated by genetic analyses [80]. Consistent with this model, loss-of-function mutations in *egl-9* or *rhy-1* and *swan-1;vhl-1* double mutations cause an array of similar phenotypes, including egg-laying defects, reduced brood size and resistance to *P. aeruginosa* PAO1 [39, 50, 81, 82].

These mutants provide an opportunity to employ multiple genetic backgrounds to over-activate HIF-1 and determine the downstream effects. This also provides some insights to the molecular networks that enable animals to respond to diverse stresses. We answer these questions by comparing the genome-wide transcriptional profiles in these HIF-1 negative regulator mutants.

## Results

### Comparisons of the transcriptional phenotypes of *vhl-1(ok161), swan-1(ok267);vhl-1(ok161), egl-9(sa307)* and *rhy-1(ok1402)*

To test our model of HIF-1 regulation and to achieve a richer understanding of the consequences of persistent HIF-1 over-activation, we employed transcriptome analyses to examine the changes in gene expression of animals carrying a deletion in *vhl-1* and animals lacking both *vhl-1* and *swan-1* functions. We also examined animals carrying strong loss-of-function mutation in *egl-9* or *rhy-1*. We reasoned that if, as current models propose, *swan-1, egl-9* and *rhy-*1 acted in common pathway(s) to inhibit HIF-1 activity, then mutations of these genes would cause similar genome-wide gene expression changes.

The complete analysis results for all the probesets on the microarray were provided in S1 Table. The list of over 2000 genes (about 10% of genes detected on the microarray) that were differentially expressed in at least one of the four mutants compared to wild-type N2 animals is provided in S2 Table. To further investigate the quality of these datasets, we compared the analysis results to earlier verified gene expressions in these mutants. By RNA blot assays, we and others had demonstrated that *nhr-57, cysl-2*/K10H10.2, F22B5.4, *rhy-1*/W07A12.7, *phy-2, fmo-2/fmo-*12, *cyp-36A1* and *egl-9* were up-regulated in mutants lacking *vhl-1* or *egl-9* function compared to N2 [41, 83, 84]. And by real-time qRT-PCR, *cysl-2*/K10H10.2 and F22B5.4 had been shown to be up-regulated in *swan-1;vhl-1* and *rhy-1* mutants compared to N2 [39, 50], and *cyp-36A1*, *clec-60*, *clec-52* were up-regulated in *egl-9(sa307)* compared to N2 [38, 41]. Consistent with these results, in our microarray experiment, these genes were up-regulated in the four HIF-1 negative regulator mutants compared to N2 (S2 Table). The one exception was expected: *rhy-1* mRNA was not over-expressed in the *rhy-1(ok1402)* deletion mutants.

Additionally, by real-time qRT-PCR, *lys-5* and *cyp-34A4* had been shown to be down-regulated in *egl-9(sa307)* compared to N2 [38]. In agreement with this, our microarray experiment showed that *lys-5* was down-regulated in *egl-9(sa307)*, *rhy-1(ok1402)* and *swan-1(ok267);vhl-1(ok161)*; and *cyp-34A4* was down-regulated in *egl-9(sa307)* and *swan-1(ok267);vhl-1(ok161)*. In addition, recently, other groups employed RNA-seq to examine the transcriptome profiles of *egl-9(sa307)*, *rhy-1(ok1402)* and *vhl-1(ok161)* [80]. Although different techniques and different statistical cutoffs were used to call differentially expressed genes, the differentially expressed genes identified by RNA-seq (fold change ≠ 1, *q*-value < 0.1) and our microarray experiment (fold change ⩾ 1.6, *q*-value ⩽ 0.05) overlap significantly: the genes identified as differentially expressed genes in *egl-9(sa307)*, *rhy-1(ok1402)* and *vhl-1(ok161)* mutants in the studies described here overlap with the RNA-seq findings by 38%, 39% and 25%, respectively (S3 Table). Taken together, the consistency between our data and prior or concurrent studies were encouraging.

We next focused on the comparisons of gene expression patterns in the three mutants with persistent HIF-1 high-activity: *egl-9(sa307)*, *rhy-1(ok1402)* and *swan-1(ok267);vhl-1(ok161)*. The model that *swan-1, egl-9* and *rhy-*1 acted in common pathway(s) to inhibit HIF-1 activity predicted that the gene expression patterns be similar and would reveal genes that were regulated by HIF-1. The overlaps of up-regulated and down-regulated gene sets were analyzed separately. We found 616 genes were up-regulated in *swan-1(ok267);vhl-1(ok161)* (S4 Table), 625 genes were up-regulated in *egl-9(sa307)* (S5 Table), and 325 genes were up-regulated in *rhy-1(ok1402)* (S6 Table). For the pair-wised overlap between *swan-1(ok267);vhl-1(ok161)* and *egl-9(sa307)*, 406 genes were co-upregulated, which accounted for 66% (= 406/616) of genes up-regulated in *swan-1(ok267);vhl-1(ok161)*, and 65% (= 406/625) of genes up-regulated in *egl-9(sa307)*. For the pair-wised overlap between *rhy-1(ok1402)* and *swan-1(ok267);vhl-1(ok161)*, 237 genes were co-upregulated, which was equal to 73% (= 237/325) of genes up-regulated in *rhy-1(ok1402)*, and 38% (= 237/616) of genes up-regulated in *swan-1(ok267);vhl-1(ok161)*. And for the pair-wised overlap between *rhy-1(ok1402)* and *egl-9(sa307)*, 265 genes were co-upregulated, this was equal to 82% (= 265/325) of genes up-regulated in *rhy-1(ok1402)*, and 42% (= 265/625) of genes up-regulated in *egl-9(sa307)*. Finally, 212 genes were commonly up-regulated in all the three HIF-1 high-activity mutants. This was equal to 34% (= 212/616 for *swan-1(ok267);vhl-1(ok161)*, 212/625 for *egl-9(sa307*)) of genes up-regulated in *swan-1(ok267);vhl-1(ok161)* or *egl-9(sa307)*, and 65% (= 212/325) of genes up-regulated in *rhy-1(ok1402)* (Fig 1A). In sum, the genes that were up-regulated in these three mutants overlapped extraordinarily to illuminate the consequences of long-term HIF-1 over-activation.

**Fig 1.**
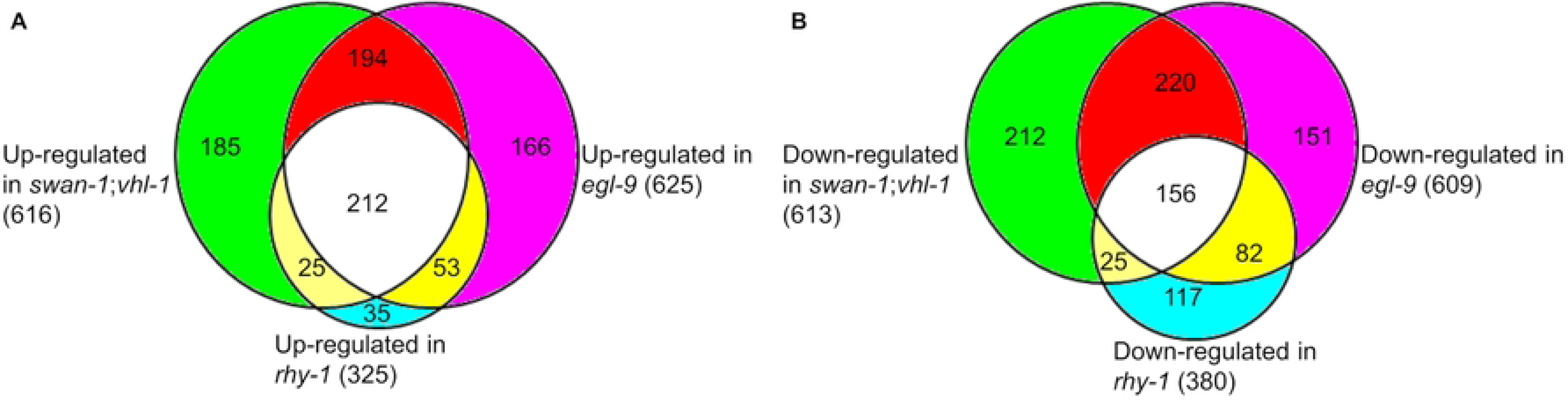
Overlaps of genes differentially expressed in the mutants with persistent HIF-1 high-activity. (A) Numbers of genes up-regulated in *rhy-1*, *egl-9* and *swan-1;vhl-1* loss-of-function mutants, relative to wild-type, and the extent to which they overlap. (B) Numbers of genes down-regulated in *rhy-1*, *egl-9* and *swan-1;vhl-1* loss-of-function mutants, and their overlaps.

We also explored the genes that were expressed at lower levels in the mutants, relative to wild-type animals. We identified 613 genes that were down-regulated in *swan-1(ok267);vhl-1(ok161)* (S7 Table), 609 genes were down-regulated in *egl-9(sa307)* (S8 Table), and 380 genes were down-regulated in *rhy-1(ok1402)* (S9 Table). For the pair-wised overlap between *swan-1(ok267);vhl-1(ok161)* and *egl-9(sa307)*, 377 genes were co-downregulated; this was equal to 62% (= 377/613) of genes down-regulated in *swan-1(ok267);vhl-1(ok161)*, and 62% (= 377/609) of genes down-regulated in *egl-9(sa307)*. For the pair-wised overlap between *rhy-1(ok1402)* and *swan-1(ok267);vhl-1(ok161)*, 239 genes were co-downregulated, this accounted for 63% (= 239/380) of genes down-regulated in *rhy-1(ok1402)*, and 39% (= 239/613) of genes down-regulated in *swan-1(ok267);vhl-1(ok161)*. For the pair-wised overlap between *rhy-1(ok1402)* and *egl-9(sa307)*, 181 genes were co-downregulated, this accounted for 48% (= 181/380) of genes down-regulated in *rhy-1(ok1402)*, and 30% (= 181/609) of genes down-regulated in *egl-9(sa307)*. Finally, 156 genes were commonly down-regulated in all the three HIF-1 high-activity mutants. This represented 25% (=156/613) of genes down-regulated in *swan-1(ok267);vhl-1(ok161)*, 26% (=156/609) of genes down-regulated in *egl-9(sa307)*, and 41% (= 156/380) of genes down-regulated in *rhy-1(ok1402)* (Fig 1B). Collectively, these analyses reveal there is a common suite of genes that are expressed at lower levels in these mutants, relative to wild-type, further defining the consequences of HIF-1 over-activation.

In sum, the microarray data revealed that the gene expression patterns in the three HIF-1 high-activity mutants were strikingly similar. This observation supported existing models that RHY-1, SWAN-1 and EGL-9 functioned in common pathway(s) to regulate HIF-1 activity.

### Consequences of persistent HIF-1 over-activation

To more fully understand the consequences of persistent HIF-1 over-activation, we explored the functions the 212 genes that were up-regulated in *swan-1(ok267);vhl-1(ok161)*, *egl-9(sa307)* and *rhy-1(ok1402)* (S10 Table), and the 156 genes that were down-regulated in these three HIF-1 high-activity mutants (S11 Table). Some biological functions were over-represented among the genes over-expressed in the mutants, and they were: carbon metabolism, biosynthesis of amino acids, sulfur metabolism, glycolysis/gluconeogenesis, innate immune response, cell division, female gamete generation, sexual reproduction, embryonic pattern specification, and response to hypoxia (Table 1). The genes that were expressed at lower levels in the mutant backgrounds, relative to wild-type, were enriched for genes with functions in organonitrogen compound catabolic process, and organic acid metabolic process (Table 2).

**Table 1.** Enriched biological terms for genes commonly up-regulated in the three mutants with persistent HIF-1 high activity.

| Biological term | Count | % | *p*-value | Genes |
| --- | --- | --- | --- | --- |
| Carbon metabolism | 10 | 5 | 3.65E-07 | *aldo-1*, C31C9.2, *cysl-1*, *cysl-2*, F26H9.5, *mce-1*, *mmcm-1*, *pcca-1*, *pccb-1*, Y62E10A.13 |
| Biosynthesis of amino acids | 8 | 4 | 3.31E-06 | *aldo-1*, C31C9.2, *cysl-1*, *cysl-2*, F26H9.5, *gln-5*, *gln-6*, Y62E10A.13 |
| Sulfur metabolism | 4 | 2 | 2.02E-04 | *cysl-1*, *cysl-2*, *ethe-1*, *sqrd-1* |
| Glycolysis/Gluconeogenesis | 4 | 2 | 6.96E-03 | *aldo-1*, D2063.1, *pck-1*, *sodh-1* |
| Innate immune response | 11 | 5 | 6.64E-03 | C14C6.5, C32H11.4, *clec-67*, *cnc-4*, *dod-24*, F08G5.6/*irg-4*, F35E12.5/*irg-5*, F55G11.8, *lys-2*, *nhr-57*, T24B8.5/*sysm-1* |
| Cell division | 6 | 3 | 5.81E-03 | *cdc-25.3*, *cpg-1*, *cpg-2*, *cyb-3*, *mei-2*, *pzf-1*, *rmd-1* |
| Female gamete generation | 14 | 7 | 6.68E-05 | *cpg-1*, *cpg-2*, *cyb-3*, *egg-1*, *egg-3*, *egg-4/egg-5*, *egg-6*, *gld-3*, *meg-1*, *nos-2*, *oma-1*, *oma-2*, *perm-1*, *rmd-1* |
| Sexual reproduction | 18 | 9 | 4.81E-03 | *cpg-1, cpg-2, cyb-3, egg-1, egg-2, egg-3, egg-4/egg-5, egg-6, gld-3, hrde-1, meg-1, nos-2, oma-1, oma-2, perm-1, rmd-1, sax-2, swm-1* |
| Embryonic pattern specification | 4 | 2 | 7.44E-03 | *meg-1, mex-1, mex-3, pos-1* |
| Response to hypoxia | 3 | 1 | 9.02E-03 | *cysl-1, egl-9, nhr-57* |

**Table 2.** Enriched biological terms for genes commonly down-regulated in the three mutants with persistent HIF-1 high activity.

| Biological term | Count | % | p-value | Genes |
| --- | --- | --- | --- | --- |
| Organonitrogen compound catabolic process | 8 | 5 | 9.87E-05 | K01C8.1, *cdd-1, ltah-1.2, glna-2, lys-5, tdo-2, gcsh-1, chil-23* |
| Organic acid metabolic process | 12 | 8 | 1.78E-04 | *ugt-46, K01C8.1, glna-2, tdo-2, acs-2, acdh-2, cth-1, ugt-30, gcsh-1, ugt-53, fat-7, odc-1* |

Since HIF complexes have been shown to regulate the expression of genes involved in glycolysis and gluconeogenesis, we anticipated that genes with these functions would be misregulated in the mutants with constitutive HIF-1 high activity. As shown in Table 1, genes enriched in the glycolysis/gluconeogenesis group included *aldo-*1 (fructose-1,6-bisphosphate aldolase), *pck-1* (phosphoenolpyruvate carboxykinase), and others.

Constitutive HIF-1 high activity also changed the expression of lipid metabolism genes, in agreement with prior studies showing that HIF-1 regulated lipid metabolism in *C. elegans* [85, 86]. These genes included *mce-1*, *mmcm-1*, *pcca-*1 and *pccb-*1 (Table 1), and *fat-7*, *acs-2* and *acdh-2* (Table 2).

*C. elegans* HIF-1 has been shown to regulate proteostasis [36, 42–44]. In our analyses, amino acid metabolism related functions were enriched. Cysteine synthase genes (*cysl-1* and *cysl-2*) and glutamine synthetase genes (*gln-5* and *gln-6*) in the biosynthesis of amino acids group were up-regulated by persistent HIF-1 high activity (Table 1). K01C8.1, *glna-2*, *tdo-2* and *gcsh-1* for amino acid catabolism in the organonitrogen compound catabolic process group were down-regulated by persistent HIF-1 high activity (Table 2).

Table 1 also includes genes involved in H2S or HCN detoxification: *cysl-1*, *cysl-2*, *ethe-1* and *sqrd-*1. This is consistent with published findings that HIF-1 was required for *C. elegans* to survive in hydrogen sulfide (H_2_S) and hydrogen cyanide (HCN) [29–34].

The finding that genes with roles in innate immune response are enriched in mutants that over-activate HIF-1 is consistent with other published studies. HIF-1 has been shown to contribute to pathogen response through regulation of C-type lectin genes, lysozyme genes and *nhr-57* [36, 38]. The transcriptome studies presented here also identify additional representative genes in this group, including caenacin antimicrobial peptide *cnc-4*, C-type lectin *clec-67,* downstream Of DAF-16 (regulated by DAF-16) *dod-24*, nuclear hormone receptor family member *nhr-57* and lysozyme *lys-2*.

It was especially interesting to see that development and reproduction related biological terms (cell division, female gamete generation, sexual reproduction, and embryonic pattern specification) were enriched for genes up-regulated by persistent HIF-1 over-activation (Table 1), as it had been shown that fecundity was decreased in *rhy-1*, *egl-9*, or *swan-1;vhl-1* mutants [39, 50]. Genes in these functional groups had roles in cell cycle progression (*cdc-25.3* and *cyb-3*), chromosome segregation (*pzf-*1 and *rmd-1*), spindle organization and rotation (*mei-2*), germline development (*gld-3)*, female gamete generation (*egg-1*, *egg-2, egg-3*, *egg-4/egg-5* and *egg-6*), and P granule segregation (*meg-1*, *mex-1* and *mex-3*) (Table 1).

### Convergence of HIF-1 pathway and pathogen immune response pathways

It had been shown that r*hy-1*(*ok1402*), *egl-9*(*sa307)* and *swan-1(ok267);vhl-1(ok161)* mutants were resistant to *P. aeruginosa* PAO1; and HIF-1 was required for defense against *P. aeruginosa* PA14 and PAO1 [36, 37, 39, 41, 81]. It also had been shown that NSY-1/SEK-1/PMK-1 mitogen-activated protein kinase pathway mediated the response to *P. aeruginosa* PA14 in *C. elegans* [87, 88]. To ask whether these two *P. aeruginosa* protective pathways included similar targets or not, we compared our datasets with published microarray studies that identified the targets of PMK-1 and SEK-1 [87].

Among the 85 genes up-regulated by PMK-1, 38 genes (45%) were co-upregulated in *swan-1(ok267);vhl-1(ok161)*, and 28 genes (33%) were co-upregulated in *egl-9*(*sa307)* (S12 Table and Fig 2A). These overlaps were significant (*p-*values were 4.44E-34 and 6.23E-21 respectively, by Fisher’s exact tests). Genes positively regulated by PMK-1 and HIF-1 included those for immune response, like lysozyme gene *lys-2*, C-type lectin genes (*clec-67* and *clec-85*), ShK-like toxin gene T24B8.5, CUB like domain gene *cld-9*, and hypersensitive to pore-forming toxin gene *hpo-6* (S12 Table).

**Fig 2.**
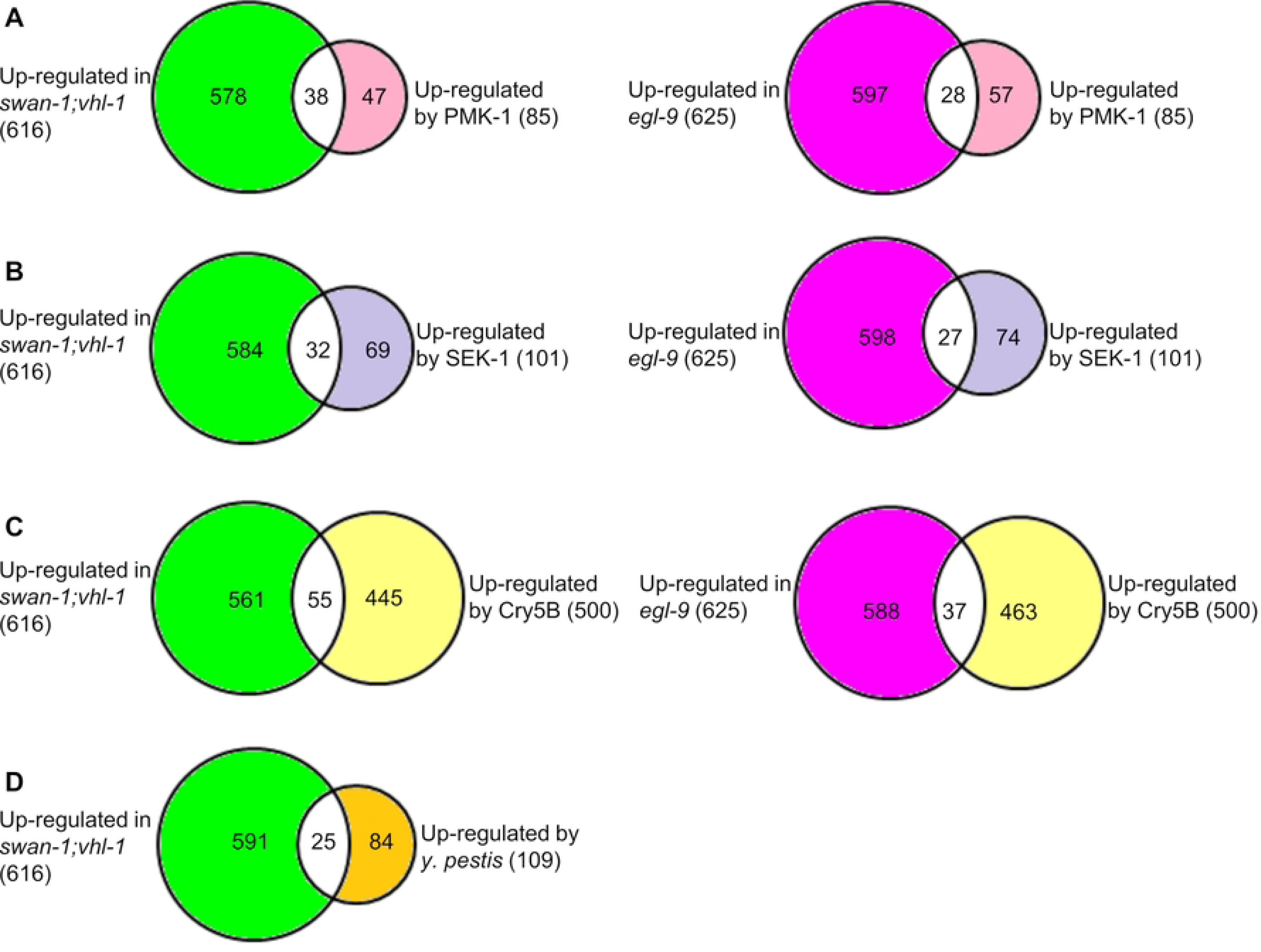
Common changes in gene expression caused by over-activation of the HIF-1 pathway and mutations in pathogen responsive pathways. (A) Genes up-regulated by PMK-1 overlapped with genes up-regulated in *swan-1;vhl-1* or *egl-9* mutants. (B) Genes up-regulated by SEK-1 overlapped with genes up-regulated in *swan-1;vhl-1* or *egl-9* mutants. (C) Genes induced by Cry5B overlapped with genes up-regulated in *swan-1;vhl-1* or *egl-9* mutants. (D) Genes induced by *Y. pestis* infection overlapped with genes up-regulated in *swan-1;vhl-1* mutants.

Genes up-regulated by HIF-1 over-activation also overlapped with genes up-regulated by SEK-1. Among the 101 genes up-regulated by SEK-1, 32 genes (32%) were co-upregulated in *swan-1(ok267);vhl-1(ok161)*, and 27 genes (27%) were co-upregulated in *egl-9*(*sa307)* (S13 Table and Fig 2B). These overlaps were significant (*p-*values were 2.30E-22 and 5.49E-16 respectively, by Fisher’s exact tests). The overlapped genes included those implicated in immunity and detoxification, such as CUB domain protein genes (*dct-17*, *cld-9* and others), C-type lectin genes (*clec-41, clec-66* and *clec-67*), UDP-glucuronosyl transferase *ugt-44,* hypersensitive to poreforming toxin gene *hpo-6*, and ShK-like toxin gene T24B8.5 (S13 Table).

We also expected to find overlaps between genes induced by HIF-1 over-activation and those activated by the crystal pore-forming toxin Cry5B, as it had been shown that *egl-9* loss-of-function mutants were resistant to Cry5B in a HIF-1-dependent manner [36]. To address this, we compared our datasets with microarray studies that identified Cry5B-responsive genes [89].

Among the 500 genes up-regulated by Cry5B, 55 genes (11%) were co-upregulated in *swan-1(ok267);vhl-1(ok161)*, and 37 genes (7%) were co-upregulated in *egl-9*(*sa307)* (S14 Table and Fig 2C). These overlaps were significant (*p-*values were 1.54E-14 and 3.10E-5 respectively, by Fisher’s exact tests). Genes responsive to HIF-1 high-activity and Cry5B included immune responsive genes, such as C-type lectin gene *clec-4*, CUB-like domain genes (*cld-9* and others), cytochrome P450 family gene *cyp-33C8*, alcohol dehydrogenase gene *sodh-1*, hypersensitive to pore-forming toxin gene *hpo-6*, and UDP-glucuronosyl transferase gene *ugt-44* (S14 Table).

Recognizing that some of the gene expression changes caused by HIF-1 over-activation were also associated with *Yersinia pestis* response [90], we asked whether genes regulated in the HIF-1 high-activity mutants overlapped with those responsive to *Y. pestis*. Among the 109 genes up-regulated by *Y. pestis*, 25 genes (23%) were co-upregulated in *swan-1(ok267);vhl-1(ok161)* (S15 Table and Fig 2D). The overlap was significant (*p-*value = 2.1E-15, by Fisher’s exact test).

Genes co-upregulated by *Y. pestis* and *swan-1(ok267);vhl-1(ok161)* double mutations included immune responsive genes, like C-type lectin genes (*clec-60*, *clec-66* and *clec-67*), alcohol dehydrogenase gene *sodh-1*, hypersensitive to pore-forming toxin gene *hpo-6*, and UDP-glucuronosyl transferase gene *ugt-54* (S15 Table).

### HIF-1 and DAF-16 function synergistically in hypoxia adaptation

We next asked whether the microarray data could illuminate the interaction between HIF-1 and DAF-16. DAF-16 is a forkhead family DNA-binding transcription factor, and it was inhibited by the sole *C. elegans* insulin-like receptor DAF-2. In *C. elegans*, DAF-16 and HIF-1 both have important roles in metabolism, stress response and longevity [35, 64, 69, 73, 87, 91–97].

However, the mechanism by which these two stress-responsive transcription factors interacted were not well understood. Therefore, it was intriguing to find that *daf-18*’s mRNA levels were up-regulated in the three HIF-1 high-activity mutants. The up-regulated fold changes were 1.8, 2.2 and 2.0 respectively in *swan-1(ok267);vhl-1(ok161)*, *egl-9(sa307)* and *rhy-1(ok1402)* compared to N2 (*q*-values were all < 0.05) (S2 Table and Fig 3A). DAF-18 is homologous to human tumor suppressor PTEN and is an important positive regulator of DAF-16 [98].

**Fig 3.**
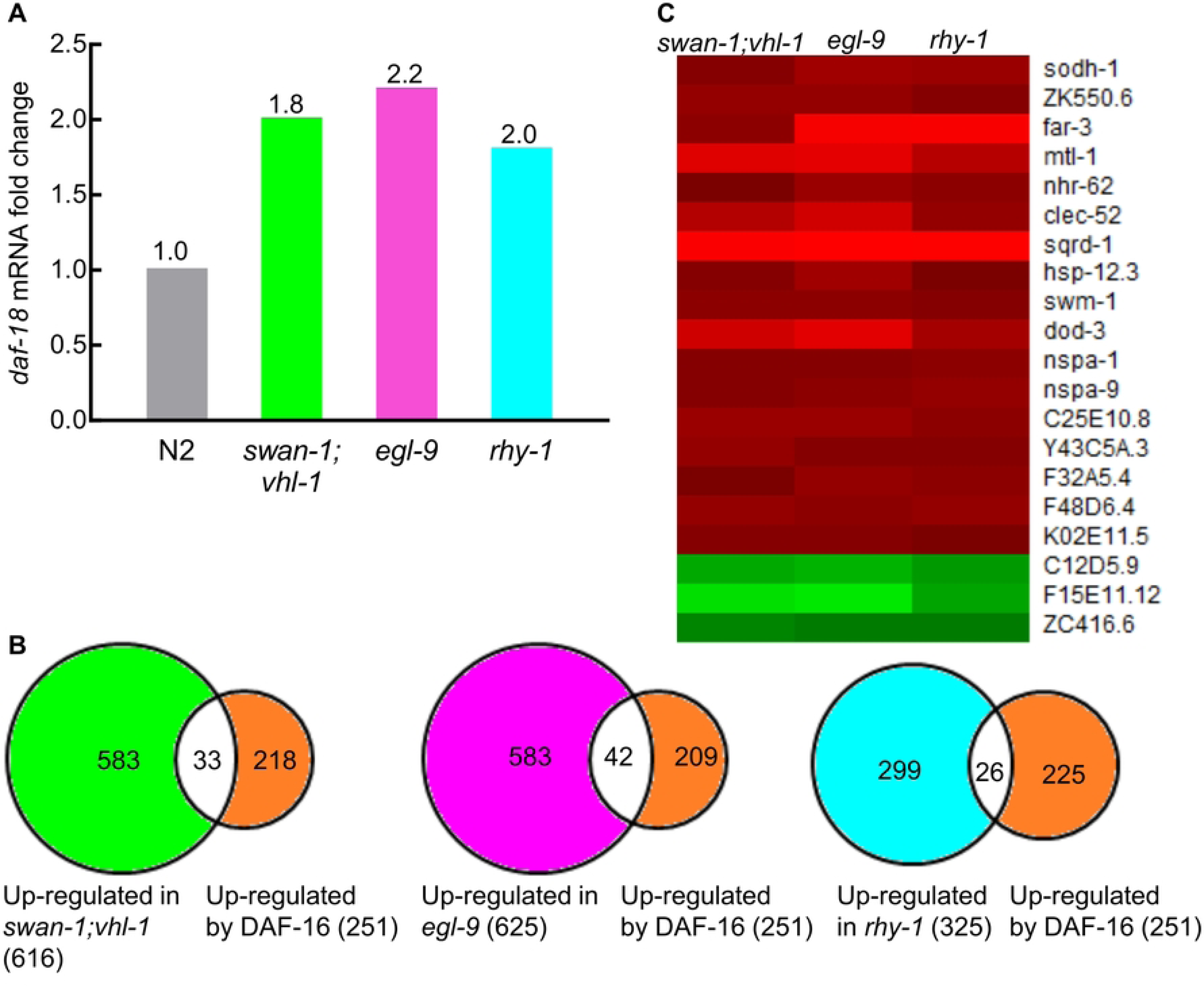
Convergence of the HIF-1 and DAF-16 pathways. (A) *daf-18*, the positive regulator of DAF-16, was up-regulated in *swan-1;vhl-1*, *egl-9* and r*hy-1* mutants. (B) Comparison of genes previously found to be up-regulated by DAF-16 [96] with genes up-regulated in *swan-1;vhl-1*, *egl-9* or r*hy-1* mutants. (C) A heat map illustrates the expression of 20 genes in *swan-1;vhl-1*, *egl-9* and r*hy-1* mutants. These genes were commonly regulated by DAF-16 and in all of these three HIF-1 high-activity mutants. The up-regulated values were represented by red color, down-regulated values were represented by green color. The color intensities correspond to the magnitudes of fold changes relative to N2 as provided in S2 Table.

We compared genes regulated by DAF-16 [96] with those regulated by HIF-1 over-activation in *swan-1(ok267);vhl-1(ok161)*, *egl-9(sa307)* or *rhy-1(ok1402)* to see whether they had significant overlaps or not. Among the 251 genes up-regulated by DAF-16, 33 genes (13%) were co-upregulated in *swan-1(ok267);vhl-1(ok161)*, 42 genes (17%) were co-upregulated in *egl-9(sa307)*, and 26 genes (10%) were co-upregulated in *rhy-1(ok1402)* (Fig 3B). These overlaps were significant, the *p*-values were 5.47E-11, 5.26E-17 and 1.42E-12, respectively, by Fisher’s exact tests.

We further identified 20 genes that were commonly regulated by DAF-16 and in all the three HIF-1 high-activity mutants. The expression profiles of these 20 genes in *swan-1(ok267);vhl-1(ok161)*, *egl-9(sa307)* and *rhy-1(ok1402)* were illustrated in a heat map in Fig 3C. This common molecular signature for DAF-16 and HIF-1 activation included increased expressions of stress-responsive and detoxification genes, such as metallothionein gene *mtl-1*, alcohol dehydrogenase gene *sodh-1*, sulfide:quinone reductase gene *sqrd-1*, small heat shock protein gene *hsp-12.3*, secreted protease inhibitor genes (*swm-1* and C25E10.8), C-type lectin gene *clec-52*, and others (Fig 3C).

We further investigated the requirement for *daf-16* in moderate hypoxia adaptation (0.5% oxygen, 21°C). Consistent with prior studies, we found that 100% of N2 eggs hatched within 24 hours in hypoxia, and 100% developed to adulthood within 72 hours. By contrast, only 52% of *hif-1(ia04)* mutant eggs hatched within 24 hours in hypoxia, and only 10% developed to adulthood within 72 hours. As shown in Table 3, loss-of-function mutations in *daf-16* also impaired embryogenesis and larval development in hypoxia. For the two *daf-16* loss-of-function alleles (*daf-16(mu86)* and *daf-16 (mgDf50)*) tested, 43-46% of *daf-16*-deficient eggs hatched within 24 hours in hypoxia, and 30-39% developed to adulthood within 72 hours (Table 3). The double mutants of *daf-16* and *hif-1* (*daf-16(mgDf50);hif-1(ia04)*) were most severely affected by hypoxia, only 8% of eggs hatched within 24 hours in hypoxia, and only 0.4% developed to adulthood within 72 hours (Table 3). Note, in room air, the hatched or adult rates for all of these mutant genotypes were the same as those in N2 wild type: 100% of the eggs hatched within 24 hours, and 100% of them developed to adulthood within 72 hours (Table 3). Using the standard provided by Kirienko et al. [99], these data suggest that HIF-1 and DAF-16 acted synergistically to protect *C. elegans* from hypoxia insult.

**Table 3.**
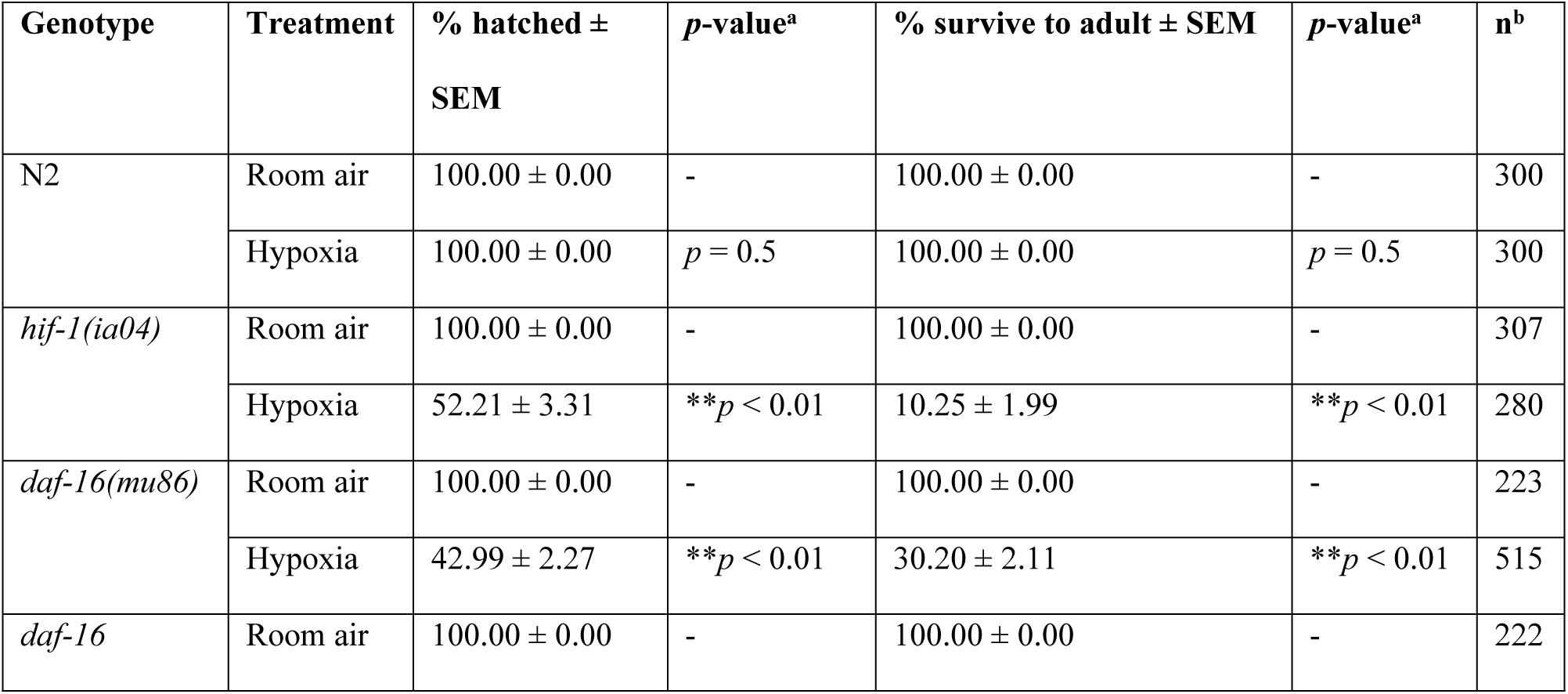
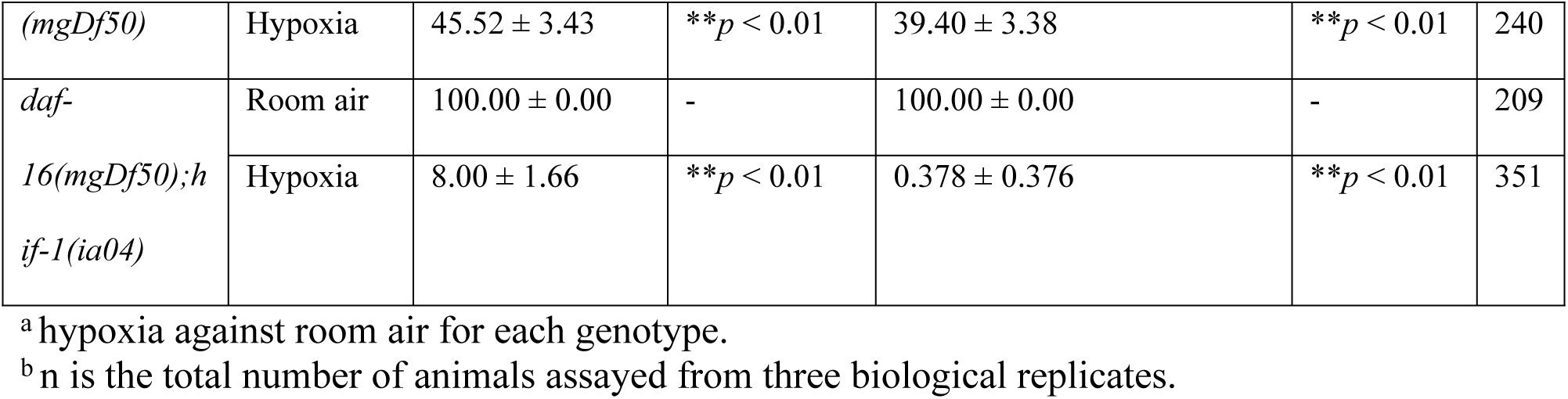
*hif-1* and *daf-16* functioned synergistically to protect *C. elegans* in hypoxia.

## Discussion

Prior studies have shown that RHY-1, EGL-9 and SWAN-1 regulate HIF-1 activity [39, 50], but the extent to which their functions overlap was not fully understood. Here, by employing whole-genome transcriptome analyses, we are able to shed more light on this important regulatory network. The analyses of genes that were differentially expressed in *rhy-1*, *egl-9* and *swan-1;vhl-1* mutants also extend our understanding of the consequences of multi-generational HIF-1 activation and provides insights to how HIF-1 pathway interacts with other pathways that mediate stress responses.

### Genome-wide gene expression analyses support RHY-1, EGL-9 and SWAN-1 function in common pathway(s) to regulate HIF-1 activity

Prior assays had shown that selected HIF-1-regulated genes were over-expressed in *egl-9*, *rhy-1* and *swan-1;vhl-1* mutants. Further, these mutants displayed similar phenotypes in terms of egg-laying defects, reduced brood size and resistance to *P. aeruginosa* PAO1 fast killing [39, 50, 81, 82]. These studies indicated that RHY-1, EGL-9 and SWAN-1 functioned in common pathway(s) to regulate HIF-1 activity. The transcriptome analyses described here and summarized in Figure 1 fortify this model.

Prior studies also show that *vhl-1*, *egl-9* and *swan-1* have HIF-1-independent functions [38, 41, 78, 83, 100]. In agreement with this, our dataset show that each mutant genotype has its own unique gene expression signature. For example, 27% (166/625) of genes up-regulated in *egl-9(sa307)* are unique to itself and not differentially expressed in *rhy-1(ok1402)* or *swan-1(ok267);vhl-1(ok161)*. Similarly, 30% (185/616) of genes up-regulated in *swan-1(ok267);vhl-1(ok161)* are not differentially expressed in *rhy-1(ok1402)* or *egl-9(sa307)*. Also, 11% (35/325) of genes up-regulated in *rhy-1(ok1402)* are not differentially expressed in *egl-9(sa307)* or *swan-1(ok267);vhl-1(ok161)* (Fig 1A). The same conclusion applies to genes down-regulated in these mutants: 24% (= 150/609) of genes down-regulated in *egl-9(sa307)* are not differentially expressed in *rhy-1(ok1402)* or *swan-1(ok267);vhl-1(ok161)*; 35% (= 212/613) of genes down-regulated in *swan-1(ok267);vhl-1(ok161)* are not differentially expressed in *rhy-1(ok1402)* or *egl-9(sa307)*; and 31% (= 117/380) of genes down-regulated in *rhy-1(ok1402)* are not differentially expressed in *egl-9(sa307)* or *swan-1(ok267);vhl-1(ok161)* (Fig 1B).

### The pleiotropic consequences of persistent HIF-1 over-activation account for its diversified biological roles

Misregulation of HIF-1 disrupts multiple facets of *C. elegans*’ life, including hypoxia adaptation, hydrogen sulfide and hydrogen cyanide resistance, heat resistance, heavy metal toxicity tolerance, pathogen response, egg-laying defect, brood size and aging [22, 29, 30, 34–36, 38, 39, 44, 50, 61–64, 81, 82]. Our analyses provide a more complete understanding of gene expression changes that underpin these diverse phenotypes. The common set of genes misregulated in *swan-1(ok267);vhl-1(ok161)*, *egl-9(sa307)* and *rhy-1(ok1402)* are involved in the metabolism of sugars, lipids, amino acids, and sulfur. Consistent with this, it have been shown that the metabolomic profiles of various amino acids, carbohydrates, lipids and nucleotides are changed in *egl-9(sa307)* mutants [101]. They also have roles in hypoxia and innate immune response, and reproduction and development. These various biological processes regulated by persistent HIF-1 over-activation suggest models for why HIF-1 can play a variety of biological roles besides regulating hypoxia response. For example, the misregulated cell cycle and reproduction processes in *swan-1(ok267);vhl-1(ok161)*, *egl-9(sa307)* and *rhy-1(ok1402)* may correspond to their similar egg-laying defects and reduced brood sizes [39, 41, 50, 82]. And the up-regulated sulfur metabolism by HIF-1 over-activation is relevant to HIF-1’s roles in hydrogen sulfide and hydrogen cyanide resistance [29, 30, 32–34]. Also, the up-regulated innate immune response by HIF-1 over-activation explains its roles in pathogen resistance [35, 36, 38–41, 81, 99], which is further supported by the overlaps of genes regulated by HIF-1 over-activation with genes responsive to toxic Cry5B and *Y. pestis* [36, 90], and with genes regulated by PMK-1 and SEK-1, the major players for pathogenic *P. aeruginosa* PA14 resistance [88].

### HIF-1 interacts with DAF-16 to promote hypoxia adaptation

DAF-16 and HIF-1 are both important stress regulators in *C. elegans*. In this study, we demonstrate that DAF-16 and HIF-1 have complementary and overlapping roles in stress response. We find that DAF-16 and HIF-1 converge on key stress-responsive genes, and hypoxia survival assays further show that HIF-1 and DAF-16 function synergistically to promote hypoxia adaptation. Intriguingly, the mRNA levels of *daf-18*, the positive regulator of DAF-16, are up-regulated in *swan-1(ok267);vhl-1(ok161)*, *egl-9(sa307)* and *rhy-1(ok1402)* mutants. This suggests a mechanism for the crosstalk between HIF-1 and DAF-16: HIF-1 may enhance DAF-16’s function by increasing the expression of *daf-18*. While beyond the scope of the experiments reported here, future studies might explore the ways in which these two important stress-response pathways regulate each other and converge to enable the organism to adapt and survive adverse conditions.

## Materials and methods

### Strains

The wild-type *C. elegans* used in this study was N2 Bristol. The loss-of-function mutation alleles used in this study were: LGI: *daf-16(mu86)lf*, *daf-16(MgDf50)lf*; LGII: *rhy-1(ok1402)lf*; LGV: *hif-1(ia04)lf*, *egl-9(sa307)lf*, *swan-1(ok267)lf*; LGX: *vhl-1(ok161)lf*. All the worms were maintained at 21°C using the standard methods [102].

### Gene expression microarray experiment

Randomized complete block design was followed for the microarray experiment, with three biological replicates treated as three blocks. Each block included eight treatments: N2 wild type, N2 wild type with hypoxia treatment, *hif-1*(*ia04)* loss-of-function mutants*, hif-1*(*ia04)* loss-of-function mutants with hypoxia treatment, *vhl-1(ok161)* loss-of-function mutants*, rhy-1(ok1402)* loss-of-function mutants, *egl-9(sa307)* loss-of-function mutants and *swan-1(ok267);vhl-1(ok161)* loss-of-function double mutants. For each treatment, about 1,000 synchronized L4-stage larvae were pooled as one experimental unit to get sufficient RNA for hybridization. Total RNA isolation was performed using Trizol (Invitrogen) and RNeasy Mini Kit (Qiagen). RNA quality was checked with an Agilent 2100 BioAnalyzer (Agilent Technologies). The RNA integrity numbers (RINs) for all the samples used in this study were greater than 9.0. The total RNA isolated from one experimental unit was hybridized onto one Affymetrix GeneChip® C. elegans Genome array (Affymetrix, part number 900383). Probe synthesis, labeling, hybridization, washing, staining and scanning were performed by the GeneChip facility at Iowa State

University. In brief, the total RNA was synthesized to biotin-labeled aRNA using the GeneChip® 3’ IVT Express Kit (Affymetrix, part number 901229) and hybridized to the array. The arrays were washed and stained in the GeneChip® fludics station 450 and scanned with GeneChip® scanner 3000 7G. The Affymetrix® GeneChip® Command Console™ (AGCC) software was used to generate probe cell intensity data (.CEL) files. The resulting CEL files were normalized and summarized using the robust multichip average (RMA) algorithm [103] in R package (R Core Team, Vienna, Austria, 2016). An analysis of variance (ANOVA) model was then fitted to the summarized expression measures, with the block (three levels) and the treatment (eight levels) treated as fixed effect factors following the experimental design.

Residual model diagnostics identified no severe violations of the model assumptions. Linear contrasts of treatment means were tested using the general F-test. To account for multiplicities of hypothesis testing, conservative estimates of false discovery rates (FDRs) were calculated according to the *q*-value procedure of Storey and Tibshirani [104]. Differentially expressed probesets were defined as *q*-value ≤ 0.05 and fold change ≥ 1.6. Probesets were converted to genes using the Affymetrix annotation file “Celegans.na36.annot.csv”. To deal with redundancy and count the number of unique genes detected on the array, we kept one probeset per gene and one gene per probeset. In this way, the total number of unique genes detected on the array was 18, 011. For the purpose of reference, the original complete lists of gene(s) annotated to each probeset were kept in S1-S8 Tables.

In this paper, we discuss gene expression changes in the mutants with persistent HIF-1 function. HIF-1-dependent gene expression changes under hypoxia will be described in a related study entitled “Whole genome profiling of short-term hypoxia induced genes and identification of HIF-1 binding sites provide insights into HIF-1 function in *Caenorhabditis elegans* ”. The microarray raw and probeset summary data had been deposited to NCBI’s Gene Expression Omnibus, the accession number was GSE228851.

### Gene function annotation and enrichment analyses

DAVID tools (The Database for Annotation, Visualization and Integrated Discovery) [105, 106] (https://david.ncifcrf.gov) were used to annotate the enriched biological terms associated with microarray-selected genes. The enriched biological terms were at *p*-value < 0.01 with no correction.

### Heat maps

Heat maps for gene expression profiles were generated using the PermutMatrix graphical analysis program [107, 108]. Average linkage clustering was performed with the fold changes compared to N2. Green color represented negative values, and red color represented positive values. The color intensities corresponded to the magnitudes of fold changes. Other parameters were set as default.

### Gene lists overlap testing

Fisher’s exact test was performed to test whether the overlap between two gene lists was significant or not. The total number of 18, 011 genes detected on the microarray was used as the population size. The significant overlap is at *p*-value < 0.001.

### Hypoxia development and survival assays

For each genotype, the room air and hypoxia treatments were performed in parallel at 21°C. For each treatment, 20 young adults (one day after L4 molt) were used as parents to lay eggs on one NGM plate seeded with OP50 for 30 minutes. After counting the eggs laid, the plates were kept in room air or put into a sealed plexiglass chamber with constant hypoxic gas flow. Compressed air and 100% nitrogen were mixed to achieve 0.5% oxygen, and gas flow was controlled by an oxygen sensor [84]. After 24 hours, the un-hatched eggs were counted for both treatments. After that, the plates for both treatments were maintained in room air. The adult worms were counted 72 hours after the eggs had been laid. The data collection time points were set to match the development rate of N2 eggs in room air: they hatch within 24 hours and reach adulthood within 72 hours. The experiments were performed with three biological replicates. To test the effect of hypoxia on animal development and survival, the binary hatched *vs*. un-hatched or adult *vs.* non-adult data were analyzed for each genotype by fitting a generalized linear model using a logit link function with JMP 9 statistical software (SAS Institute Inc., Cary, NC, 2010). The replicate (three levels) and the treatment (two levels) were used as factors in the model.

## Acknowledgments

The experiments reported here were conducted as part of Dingxia Feng’s PhD thesis work, under the guidance of Dr. Jo Anne Powell-Coffman at Iowa State University. The work was supported by grant R01GM078424 from the National Institutes of Health to JP. Author DF conducted the experiments and drafted the manuscript, and author LQ was instrumental to the statistical design and analyses of gene expression comparisons. The authors are grateful for mentors and thesis committee members who provided advice and guidance.

## Supporting information

**S1 Table. Gene expression for all the probesets under hypoxia and in the HIF-1 negative regulator mutants.**

**S2 Table. Genes differentially expressed in *vhl-1*, *swan-1;vhl-1*, *egl-9* or *rhy-1*.**

**S3 Table. Overlaps between differentially expressed genes (DEGs) identified by RNA-seq and microarray.**

**S4 Table. Genes up-regulated in *swan-1;vhl-1*. S5 Table. Genes up-regulated in *egl-9*.**

**S6 Table. Genes up-regulated in *rhy-1*.**

**S7 Table. Genes down-regulated in *swan-1;vhl-1*. S8 Table. Genes down-regulated in *egl-9*.**

**S9 Table. Genes down-regulated in *rhy-1*.**

**S10 Table. Genes commonly up-regulated in *swan-1;vhl-1*, *egl-9* and *rhy-1*. S11 Table. Genes commonly down-regulated in *swan-1;vhl-1*, *egl-9* and *rhy-1*. S12 Table. Genes co-upregulated by PMK-1 and HIF-1.**

**S13 Table. Genes co-upregulated by SEK-1 and HIF-1. S14 Table. Genes co-upregulated by Cry5B and HIF-1.**

**S15 Table. Genes co-upregulated by *Yersinia pestis* and HIF-1.**

